# mzMLb: a future-proof raw mass spectrometry data format based on standards-compliant mzML and optimized for speed and storage requirements

**DOI:** 10.1101/2020.02.13.947218

**Authors:** Ranjeet S. Bhamber, Andris Jankevics, Eric W Deutsch, Andrew R Jones, Andrew W Dowsey

**Affiliations:** Department of Population Health Sciences and Bristol Veterinary School, University of Bristol BS8 2BN, United Kingdom; School of Biosciences and Phenome Centre Birmingham, University of Birmingham, Birmingham B15 2TT, United Kingdom; Institute for Systems Biology, Seattle, Washington 98109, United States; Institute of Integrative Biology, University of Liverpool, Liverpool L69 7ZB, United Kingdom

**Keywords:** Proteomics Standards Initiative, mzML, Mass Spectrometry, proteomics, metabolomics, data compression, HDF5

## Abstract

With ever-increasing amounts of data produced by mass spectrometry (MS) proteomics and metabolomics, and the sheer volume of samples now analyzed, the need for a common open format possessing both file size efficiency and faster read/write speeds has become paramount to drive the next generation of data analysis pipelines. The Proteomics Standards Initiative (PSI) has established a clear and precise XML representation for data interchange, mzML, receiving substantial uptake; nevertheless, storage and file access efficiency has not been the main focus. We propose an HDF5 file format ‘mzMLb’ that is optimised for both read/write speed and storage of the raw mass spectrometry data. We provide extensive validation of write speed, random read speed and storage size, demonstrating a flexible format that with or without compression is faster than all existing approaches in virtually all cases, while with compression, is comparable in size to proprietary vendor file formats. Since our approach uniquely preserves the XML encoding of the metadata, the format implicitly supports future versions of mzML and is straightforward to implement: mzMLb’s design adheres to both HDF5 and NetCDF4 standard implementations, which allows it to be easily utilised by third parties due to their widespread programming language support. A reference implementation within the established ProteoWizard toolkit is provided.

## Introduction

Through an extensive industry-wide collaborative process, in 2008 the Proteomics Standards Initiative (PSI) established a standardised XML (Extensible Markup Language) representation for raw data interchange in mass spectrometry (MS)^1^, ‘mzML’, further building upon concepts defined in earlier formats mzData and mzXML^2^ formats. mzML is now the pervasive format for interchange and deposition of raw mass spectrometry (MS) proteomics and metabolomics data^3^. However, in order to provide a detailed, flexible, consistent and simple standard for the sharing of raw MS data, it was designed around a generic ontology for its representation at the expense of inefficient storage and file access. Two data types are contained within raw mass spectrometry (MS) datasets: (a) numeric data i.e. mass over charge and spectral/chomatographic intensities; and (b) metadata related to instrument and experimental settings. mzML encodes these data types within a rich, schema-linked XML file, where the metadata is accurately and unambiguous annotated using the PSI-MS controlled vocabulary^4^ (CV). However, one of the bottlenecks of mzML’s design is that it is a text-based XML file format and all numeric spectrum data are converted into text strings using Base64 encoding^5^. Optionally, the numeric data can be zlib^6^ compressed before encoding, but nevertheless, the size of output files are still 4- to 18-fold higher than the original proprietary vendor format.

A number of technologies^7–9^ have been developed by various laboratories to address the inherent performance/practical difficulties of utilizing the mzML format for large volume biological sampled, high throughput data analysis. The first approach to address the performance and file size issues of mzML was mz5^7^. At the core of mz5 is HDF5^10^ (Hierarchical Data Format version 5), originally developed by the National Center for Supercomputing Applications (NCSA) for the storage and organization of large amounts of data. HDF5 is a binary format, but is similar to XML in the sense that files are self-describing and allow complex data relationships and dependencies. An HDF5 file allows multiple datasets to be stored within it in a hierarchical group structure akin to folders and files on a file system. The two primary objects represented in HDF5 files are ‘groups’ and ‘datasets’. Groups are container constructs that are used to hold datasets and other groups. Datasets are multidimensional arrays of data elements of a specific type *e.g.* integer, floating point, characters, strings, or a collection of these organised as compound types. Both objects support metadata in the form of attributes (key-value pairs) that can be assigned to each object; these attributes can be of any data type. Using groups, datasets and attributes, complex structures with diverse data types can be efficiently stored and accessed. Each dataset can optionally be subdivided into regular ‘chunks’ to enable more efficient data access, as chunks can be loaded and stored in HDF5’s cache implementation for subsequent repeated access. By changing the chunk size parameter, it is possible to adjust HDF5 for different applications, e.g. fast random access where file size does not matter, or larger chunks for an overall smaller compressed file size.

Compared with mzML, mz5’s implementation in HDF5 yields an average file size reduction of 54% and increases linear read and write speeds 3–4-fold^7^. However, mz5 involves a complete reimplementation of mzML accomplished through a complex mapping of mzML tags and binary data to compound HDF5 datasets that mimic tables in a relational database. This structure would need to be explicitly altered to accommodate future versions of mzML. The mapping also precludes a Java implementation using the HDF5 Java API as compound structures are extremely slow to access with this API. Moreover, some implementation choices are not supported by the Java API at all, specifically the variable-length nested compound structures mz5 uses to describe scan precursors.

The mzDB format^8^ uses an alternative database paradigm, the lightweight SQLite relational database. mzDB’s main mechanism of increasing random read performance is in organizing data in small two-dimensional blocks across multiple consecutive spectra (*i.e.* along both the m/z and retention time axis). In comparison with XML formats, mzDB saves 25% of storage space and improves access times by two-fold or more, depending on the data access pattern. Due to its unique data indexing and accessing scheme, three different software libraries have been created to handle MS datasets, two of which are designed to create and handle MS Data-dependent acquisition (DDA), the first “pwiz-mzDB” and the second, “mzDB-access”. The third instance named “mzDB-Swath” is specifically designed for the Data-Independent Acquisition (DIA) MS-SWATH technique. In order to utilize mzDB for other or future methods, these libraries will need to be extended. In addition, mzDB does not compress the text metadata, which are stored in dedicated “param_tree” fields in XML format with specific XML schema definitions (XSDs). mzDB also stores raw datasets uncompressed, but compression can be achieved through an SQLite extension; however, this extension requires a commercial license for both compression and decompression, and comparative results were not presented in the manuscript. As an alternative, a “compressed fitted” mode is proposed which centroids each peak and records the left and right Half Width at Half Maximum (LHW/RHW) for reconstruction; nevertheless, centroiding is insensitive to low-intensity and overlapping peaks, hence much information is discarded in the process, which may affect results when this loss of data is propagated through downstream analysis pipelines.

The imzML data format^11^ is predominantly aimed at storing very large mass spectrometry imaging datasets and does so through modest modifications to the mzML format. At the core of this approach is the splitting of XML metadata from the binary encoded data into separate files (*.imzML for the XML metadata and *.ibd for the binary data) and linking them unequivocally using a universally unique identifier (UUID). imzML also introduces new controlled vocabulary (CV) parameters designed specifically to facilitate the use of imaging data. These additional imzML CV parameters include x/y position, scan direction/pattern, pixel size are stored in the *.obo file, following OBO format 1.2 (which is text based format used to describe the CV terms). The approach is designed to enable easier visualisation of the data using third party software.

Unlike mz5, mzDB and imzML, Numpress^9^ is an encoding scheme for mzML and not a new or modified file format; its main focus on improving the file size is based on a novel method to compress the binary data in the mzML file before Base64 encoding (note: it does not compress the XML metadata). It accomplishes this by encoding the three common numerical data types present in mzML (mass to charge ratios – m/z, intensities and retention times) using a variety of heuristics. The first, Numpress Pic (numPic), is intended for ion count data (e.g. from Time of Flight) and simply rounds the value to the nearest integer for storage in truncation form. The second, Numpress Slof (numSlof), is for general intensity data and involves a log transformation followed by a multiplication by a scaling factor and then conversion and truncation to an integer. This ensures an approximately constant relative error; the authors demonstrate that choosing the threshold to yield a relative error of < 2×10^−4^ did not noticeably affect downstream analysis results. The third approach, Numpress Lin (numLin), is intended specifically for m/z values and uses a fixed-point representation of the value, achieved by multiplying the data by a scaling factor and rounding to the nearest integer. Likewise, a relative error of approximately < 2×10^−9^ was deemed not to unduly affect downstream processing. Taken together, Numpress was shown to reduce mzML file size by around 61%, or approximately 86% if the Numpress spectral output was additionally zlib compressed.

In the proposed mzMLb format, we adopt the HDF5 format^10^ also used by mz5, which is well-established for high-volume data applications. However, rather than using a complex and inflexible mapping between mzML and HDF5, we propose a simple hybrid format where the numeric data are stored natively in HDF5 binary while the metadata are preserved as fully PSI-standard mzML and linked to the binary in a manner similar to imzML - but stored within the same HDF5 file. Furthermore, we use only core features of HDF5, making our format compatible with NetCDF4^12^ readers and writers (including their native Java library). This enables third party bioinformatics tool developers to import and export data written in mzMLb using libraries already available on a wide variety of platforms and programming languages in a straightforward way. Taking advantage of inbuilt HDF5 functionality, we also implement a simple predictive coding method that enables efficient lossy compression that results in file sizes comparable to Numpress but is much easier to implement. Alternatively, Numpress compressed data can be stored in mzMLb without modification. We provide a reference implementation for mzMLb in the popular ProteoWizard toolset, available at https://github.com/biospi/pwiz.

## Methods

The fundamental design of mzMLb is shown in Figure 1, with the full specification given in the supplementary material. As illustrated, an mzMLb HDF5 file is composed of datasets for different data types (numerical and text based) contained within an mzML file. In this example with our ProteoWizard implementation, the data is stored in four HDF5 datasets: Chromatogram start scan times (chromatogram_MS_1000595_double); chromatogram intensities (chromatogram_MS_1000515_float); spectrum m/z’s (spectrum_MS_1000514_double); and spectrum intensities (spectrum_MS_1000515_float). These datasets are accompanied by native HDF5 version mirroring the indexed mzML schema (e.g. mzML_chromatogramIndex and mzML_chromatogramIndex_idRef). It illustrates how mzMLb utilises the advantages of mzML (XML) and propriety binary vendor formats by combining the positive values of both approaches while mitigating the negative traits.

**Figure 1.**
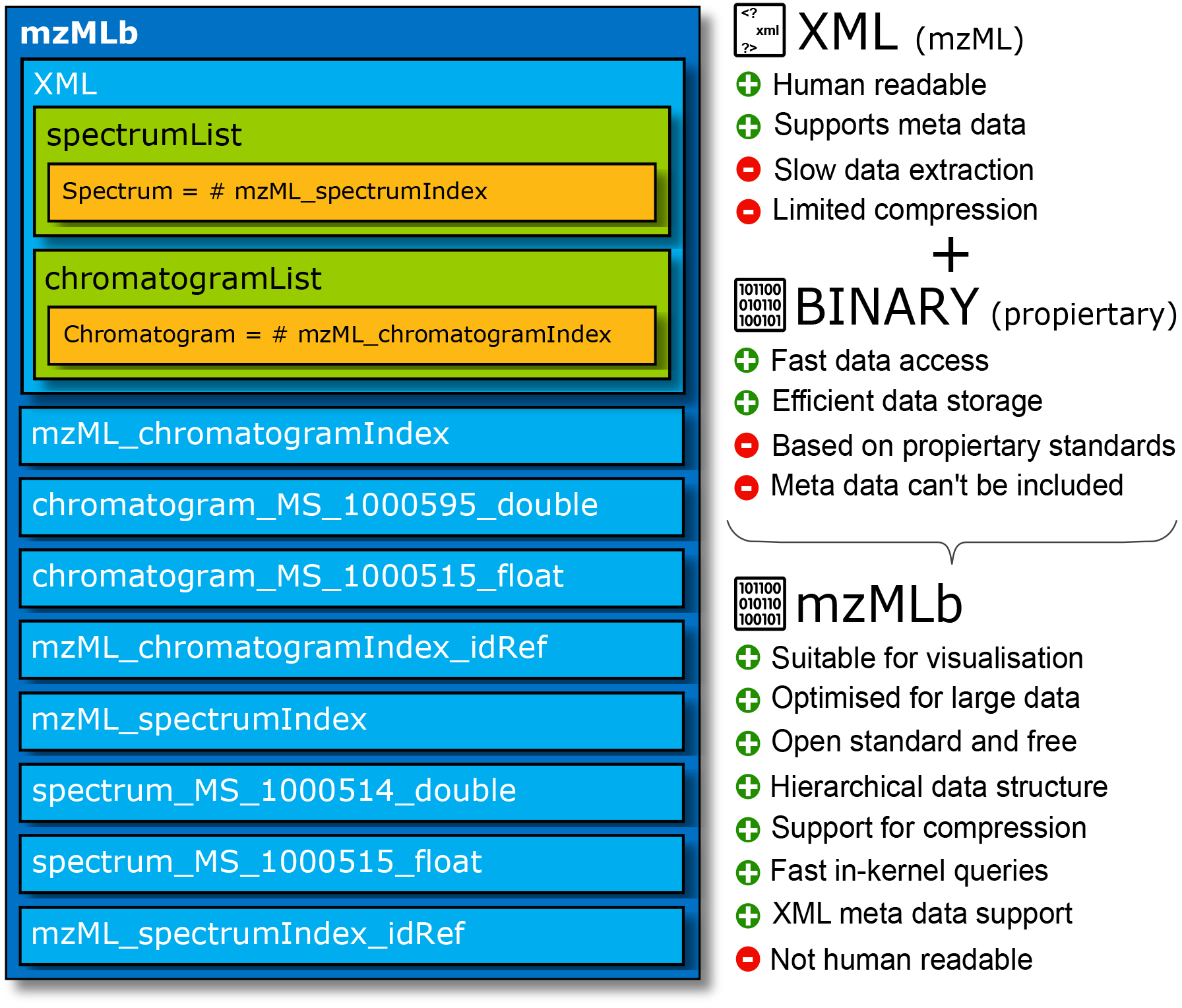
mzMLb Internal data structure. All data is stored using standard HDF5 constructs, PSI-standard mzML is maintained, full XML metadata is stored, along with binary data in separate HDF5 datasets. Storage of the chromatogram and spectral data (scan start times, m/z’s and intensities) is flexible and self-described in terms of floating-point precision and layout, relying simply on the dataset name and offset being specified within the <binary> tag for each chromatogram and spectrum in the mzML XML metadata.

The mzML XML metadata is stored inside a HDF5 character array dataset ‘mzMLb’. This is identical to the mzML format except: (a) The binary data is not stored within the <binary> tags; instead, the binary tag provides attributes for the name of the HDF5 dataset containing the binary data, and the offset within the HDF5 dataset where the data is located. This mechanism is also used in imzML, and results in valid mzML. (b) If mzML spectrum and chromatogram indices are desired (i.e. an <indexedmzML> block in mzML), they are represented instead by native HDF5 datasets ‘mzML_spectrumIndex’ and ‘mzML_chromatogramIndex’, which are one dimensional arrays of 64-bit integers pointing to the start byte of each spectrum/chromatogram in the ‘mzMLb’. In addition, spectrum/chromatogram identifiers, spot ID (an identifier for the spot from which this spectrum was derived, if a MALDI or similar run) and scan start time indices can be specified as further HDF5 datasets (see supplementary material).

All numerical data that is Base64 encoded in mzML (m/z’s, intensities, etc.) is instead stored in mzMLb as native HDF5 datasets, either as floating-point (32-bit or 64-bit), or as a generic byte array if Numpress encoded. As each <binary> tag in the ‘mzMLb’ dataset specifies the name of the dataset containing the data, each mzMLb implementation has the freedom to organise the binary data as it wishes. Since offsets can be specified, data from multiple spectra can also be co-located within the same HDF5 dataset, as long as they are of the same data type. This enables mzMLb to harness efficiency gains from HDF5 chunk-based random access and caching schemes, and also reduces file size as data will then be compressed across spectra (which is not possible in mzML). In our ProteoWizard reference implementation of mzMLb, chromatogram and spectrum data are kept apart but otherwise all data for a specific controlled vocabulary parameters (CVParam) are stored in the same dataset. For example, in the dataset in Figure 1 spectrum intensity values for all spectra are stored in dataset ‘spectrum_MS_1000515_float’.

We also implemented a simple coding scheme that combines data truncation, a linear prediction method and use of HDF5’s inbuilt ‘shuffle’ filter to improve the results of a subsequent compression step. The aim of this approach is to exploit the way numerical floating-point data is represented in binary natively on modern computing hardware, resulting in much better compression ratios. The method is lossy but like Numpress is designed only to add relative error at very small parts per million that does not affect downstream processing. Compared to Numpress, it is much easier to implement by third party developers as the encoding and decoding can be implemented in a single line of code.

In order to fully appreciate its function and implementation, a basic understanding of how decimal real numbers are represented as binary floating-point numbers is required. A number in double-precision (64-bit) or single-precision (32-bit) binary floating point^13^ format consists of three parts; a sign, an exponent and a mantissa, as represented in Figure 2. The sign bit represents a negative or positive number if set or unset respectively (blue binary bit in fig. 2). The exponent bits represent the scale of the number, and hence specifies the location of the decimal point within the number (orange binary bits in fig. 2). Finally, the mantissa (green binary bits in fig. 2) expresses the fractional part of the number - the number of bits in the mantissa hence gives you the number of significant figures. Having more bits in the exponent (11 bits in double precision compared to 8 bits in single precision) allows you to represent a wider range of numbers, whereas more bits in the mantissa (e.g. there are 52 bits in double-precision versus 23 bits in single-precision) allows more precision. If full precision is not required, then a large number of bits are stored unnecessarily, resulting in unnecessary memory and storage use. This is the case for a significant amount of the numerical data stored in conventional mzML files.

**Figure 2.**
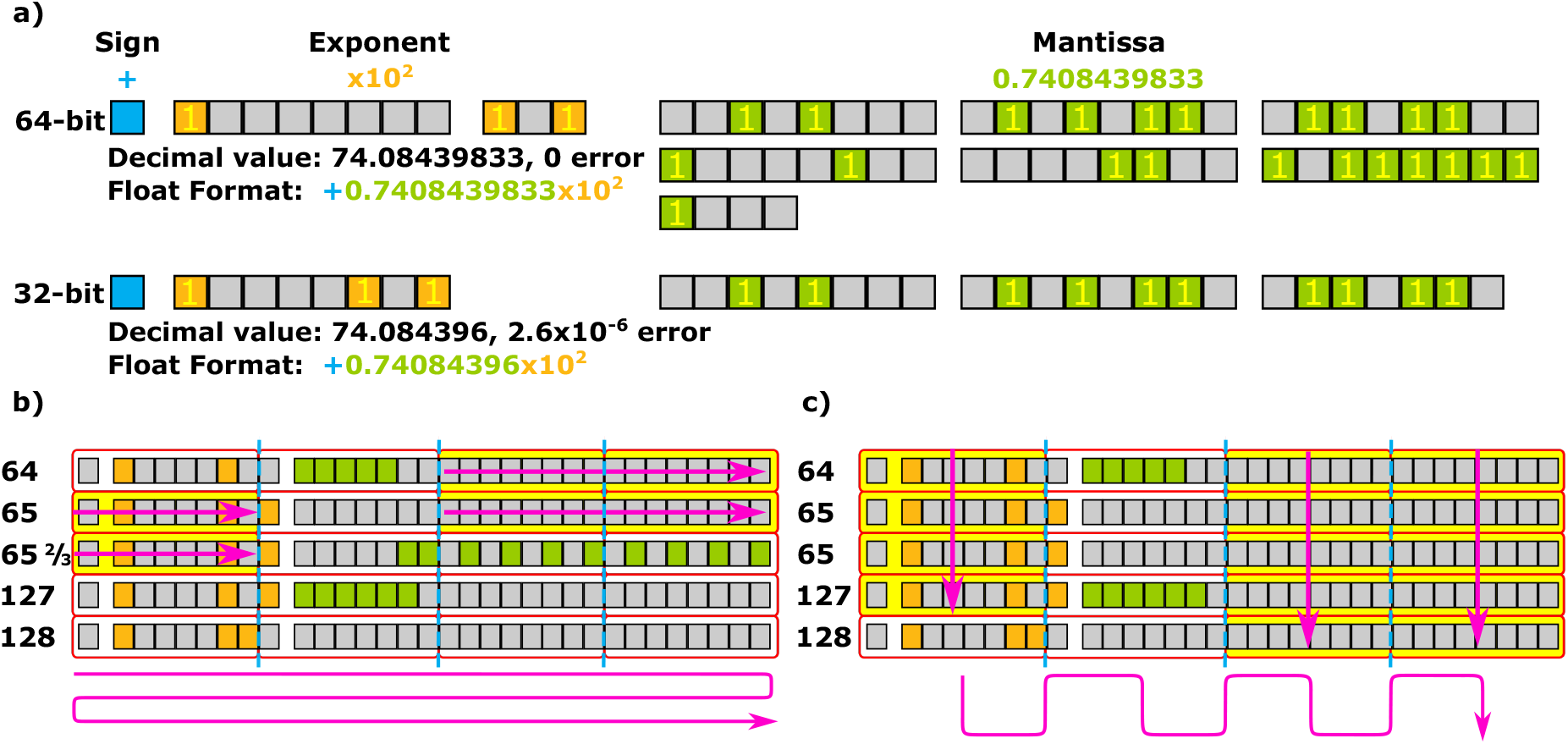
(a) Visual representation of IEEE 754 Double-precision (64 bit) floating point format and IEEE 754 Single-precision (32 bit) floating point forma, zeros are represented by empty boxes and ones are populated. (b) An array of floating point numbers stored conventionally; yellow bytes can be compressed. (c) The same array truncated and stored using the HDF5 shuffle filter leads to higher compressibility.

We exploit this fact by implementing a simple lossy truncation scheme based on reducing the numbers of mantissa bits used in the floating point format to represent m/z and intensity values by zeroing insignificant bits, with an example shown in Table 1. Here we can see that we do not observe an appreciable drop in the parts-per-million accuracy of the decimal number until after we remove 21 bits from the mantissa, and it can be seen how zeroing more and more bits increases the error as we pass the single-precision (23-bits) mantissa level.

**Table 1.** Effect of changing the number of bits representing the mantissa in a floating-point number and the associated error. The mantissa of a double-precision (64-bit) floating-point number (52 bits in the mantissa), the mantissa of a single-precision (32-bit) floating-point number (23 bits in the mantissa) are both, highlighted in green accordingly.

| Mantissa Size | Mantissa Binary | Truncated Decimal | Error Parts per million |
| --- | --- | --- | --- |
| 52 | 1001000000010101100110110010000100000011001011111110 | 400.08439833 | 0.00000 |
| 46 | 1001000000010101100110110010000100000011001011 | 400.084398329996 | 8.80887e-09 |
| 39 | 100100000001010110011011001000010000001 | 400.084398329724 | 6.90786e-07 |
| 32 | 10010000000101011001101100100001 | 400.084398329258 | 1.85469e-06 |
| 27 | 100100000001010110011011001 | 400.084398269653 | 0.00015 |
| 24 | 100100000001010110011011 | 400.084396362305 | 0.00492 |
| 23 | 10010000000101011001101 | 400.084381103516 | 0.04306 |
| 21 | 100100000001010110011 | 400.084350585938 | 0.11933 |
| 20 | 10010000000101011001 | 400.084228515625 | 0.42445 |
| 17 | 10010000000101011 | 400.083984375 | 1.03467 |
| 16 | 1001000000010101 | 400.08203125 | 5.91645 |
| 14 | 10010000000101 | 400.078125 | 15.68002 |
| 13 | 1001000000010 | 400.0625 | 54.73428 |

In order to translate our truncation approach into improved zlib^6^ compression it is necessary to employ HDF5 byte shuffling. In most formats, floating point numbers are stored consecutively on disk, so zeroed mantissa bits appear in short bursts, as shown in Figure 2b. The HDF5 shuffle filter rearranges the byte ordering of the data so that it is stored transversely rather than longitudinally, as shown in Figure 2c. This leads to large numbers of consecutive zeros that can be compressed extremely well.

Moreover, further gains are possible by transforming the data so that consecutive values or sets of values are identical, as zlib is designed to compress away repeated patterns. Towards this goal, the mz5 format uses a ‘delta’ prediction scheme that stores the difference between consecutive data points, rather than the data points themselves. This results in floating point bit patterns (Figure 2) that are less likely to change between consecutive data points and hence are more likely to be compressed. We present an improved technique termed ‘mzLinear’ that extends this approach to a linear extrapolation predicting each data point from the two previous data points, with only the error between the prediction and the actual value stored. As there is often a quadratic relationship across m/z values (for example, since there is a quadratic relationship between time-of-flight and m/z for a standard time-of-flight analyser), the aim of ‘mzLinear’ is to result in an approximately constant prediction error across the m/z range, which will compress extremely well. In comparison, delta prediction on quadratic data would lead to prediction errors that rise linearly with m/z. The technique and equation to calculate the stored error Δ*h* is depicted in Figure 3, with the plot showing a numerical series of m/z values exhibiting a quadratic relationship and how the prediction error Δ*h* remains constant for each value.

**Figure 3.**
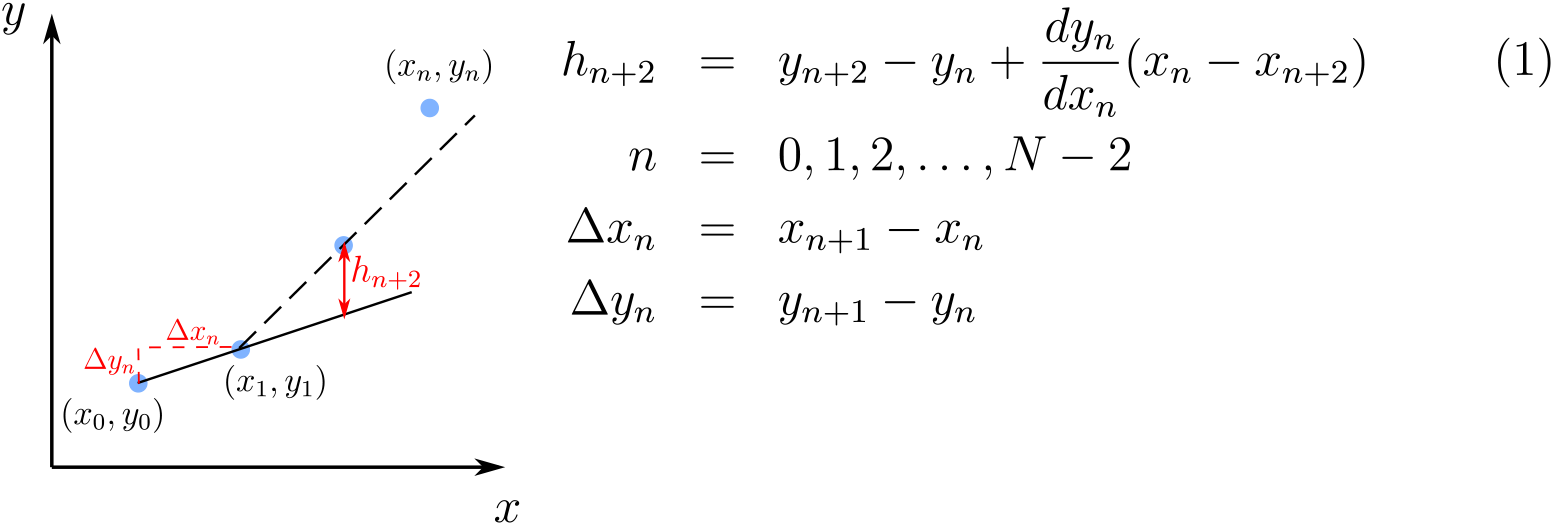
mzLinear; linear predictor implemented in mzMLb, where m/z_n_ = y_n_ and the index i_n_ = x_n_, both, *h*_0_ = 0 and *h*_1_ = 0 as the first value is stored in the new array and a linear equation can always be derived to intersect the first two points. However, for the rest of the data points; *h*_*n*+2_ where n = 0, 1, 2…, N−2, is calculated by linear predictor equation based on the previous two points and N is the total number of m/z values.

In order to demonstrate mzMLb across a broad spectrum of proteomics and metabolomics datasets used in different laboratories, we selected a wide variety MS techniques and instruments from varying vendors. The datasets are depicted in the supplementary material Table S1; files 1 through to 4 are from^9^, data files 5 to 8 are from^14^, 9 is from^15^, 10 is from^16^, 11 is from^17^ and finally 12 is from^18^. We tested mzMLb across different MS types including SWATH-DIA, DDA and Selected reaction monitoring (SRM) data, and from the major vendors including Thermo, Agilent, Sciex and Waters. Our implementation of mzMlb has been integrated into the open source cross-platform ProteoWizard software libraries and tools, and is available from https://github.com/biospi/pwiz. Hence, the proprietary raw vendors files can be directly converted into mzMLb using the ‘msconvert’ tool.

## Results

We first analyze the performance and generalisability of our truncated mzLinear coding method for m/z accuracy. Figure 4 shows the effects of the change of mantissa on the dataset ‘AgilentQToF’; it can be seen that increasing truncation decreases the file size while having minimal effect on accuracy. The effectiveness of the mzLinear prediction clearly improves zlib compression rates significantly across the range of possible truncations, as it is able to exploit the quadratic nature of the m/z to time of flight relationship.

**Figure 4.**
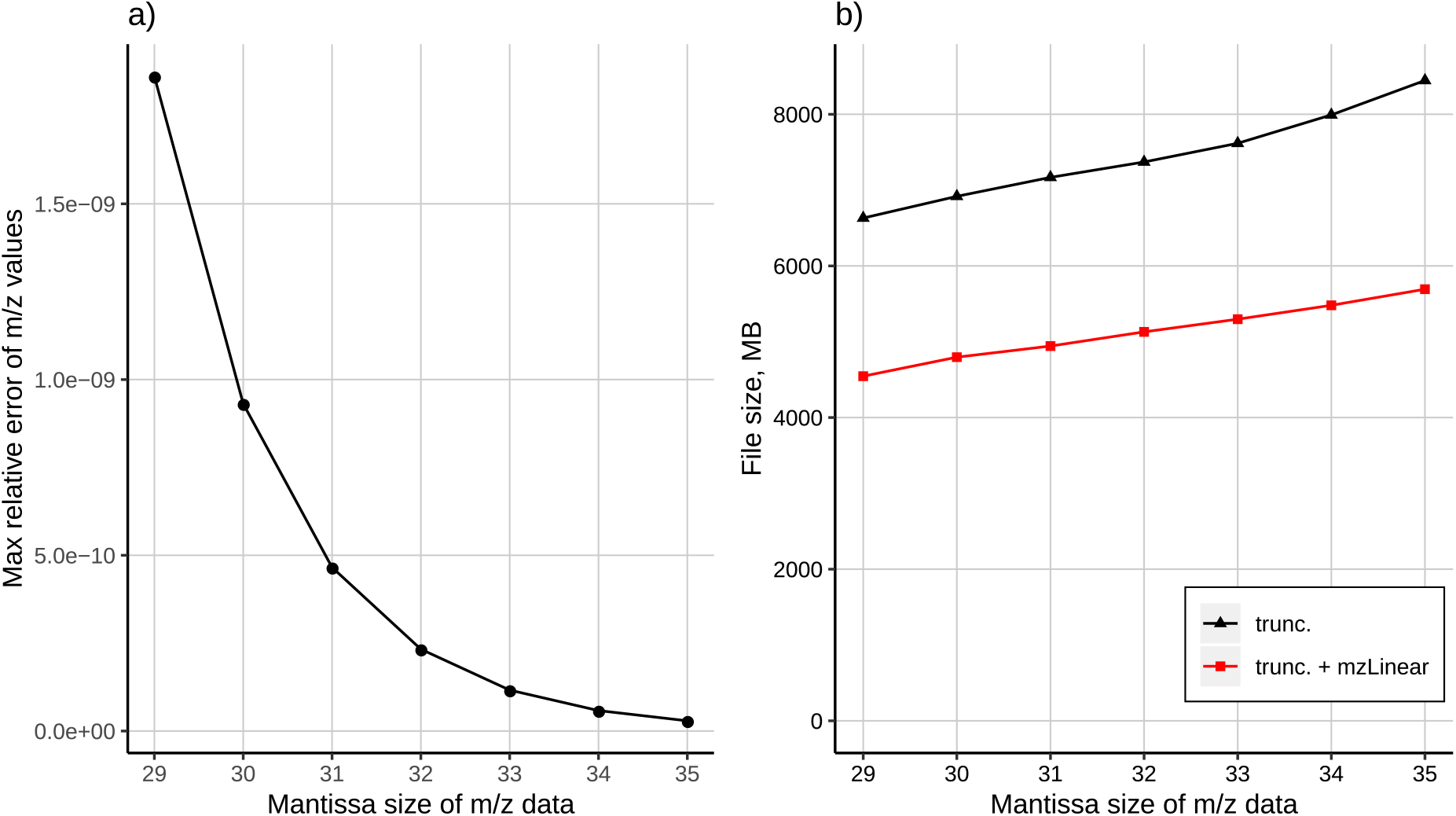
mzMLb; Mantissa truncation of AgilentQToF data file; with truncation error and file size for both mzLinear enabled and disabled.

The procedure was performed on all datasets tested, and the mantissa values were chosen such that the error induced by truncation would be less than that or comparable to Numpress’ default values of < 2×10^−4^ and < 2×10^−9^ relative errors for the intensities and m/z values respectively, which according to^9^ are small enough so as to have no effect on the output of results on the downstream of a given workflow. The result of the relative errors can be seen in the supplementary material Table S2, where mzMLb produces higher compression ratios and hence smaller files sizes.

Since mzMLb’s truncation relative error is always less than that of Numpress, the validation that Numpress does not noticeably affect downstream processing^9^ applies also to mzML. Moreover, we expand this validation by compressing the AgilentQToF and QExactive data files (shown in Table S1) and processing these files through a Mascot peptide search and protein inference workflow, the results of which can be seen in Figure 5. For the case of the AgilentQToF data we found that Numpress was unable to produce exactly the peptide and protein lists as the original uncompressed datafile. However, mzMLb was able to produce the same search and inference results for both the peptide/protein list as the original. Here we can see that the relationship between the peptide E-values of both mzMLb and Numpress against the original dataset, mzMLb gives an injective mapping (a straight line) vs the original peptide E-value, whereas the Numpress results are unable to produce the same injective relationship. The discontinuity of the Numpress results can be further illustrated by observing the peptides in the shaded regions of Figure 5c; the peptides highlighted in the enclosed box *α* represent the peptides that were present in both the original file and mzMLb but failed to be found in Numpress, whereas the peptides enclosed in the *β* region are peptides that were found in Numpress results but were not present in both the original and mzMLb results. The number of peptides deviating from the original file can be seen in Figure 5a, here we see that Numpress did not perform quite as well as mzMLb as there are a small number of peptide score values deviating from the original dataset. In Figure 5b and 5d we can see the results of the same procedure on the QExactive file, here we can see that mzMLb again produces an injective relationship with the original dataset *i.e*. producing the same results as the unmodified dataset. Numpress in Figure 5b preforms much better with an extremely low number of peptides showing ΔS deviation. However, Numpress is still unable to produce exactly the same results as the original in the Mascot pipeline. It does, however, perform better than the AgilentQToF case and demonstrates that the lossy compression method employed in Numpress is more susceptible to different vendor data files, whereas mzMLb truncation scheme is more robust to datafile vendor variation and able to reproduce the same results as an unmodified datafile. We thus take the most conservative truncation values from Table S2 (AgilentQToF truncation values) and apply them as the mzMLb defaults.

**Figure 5.**
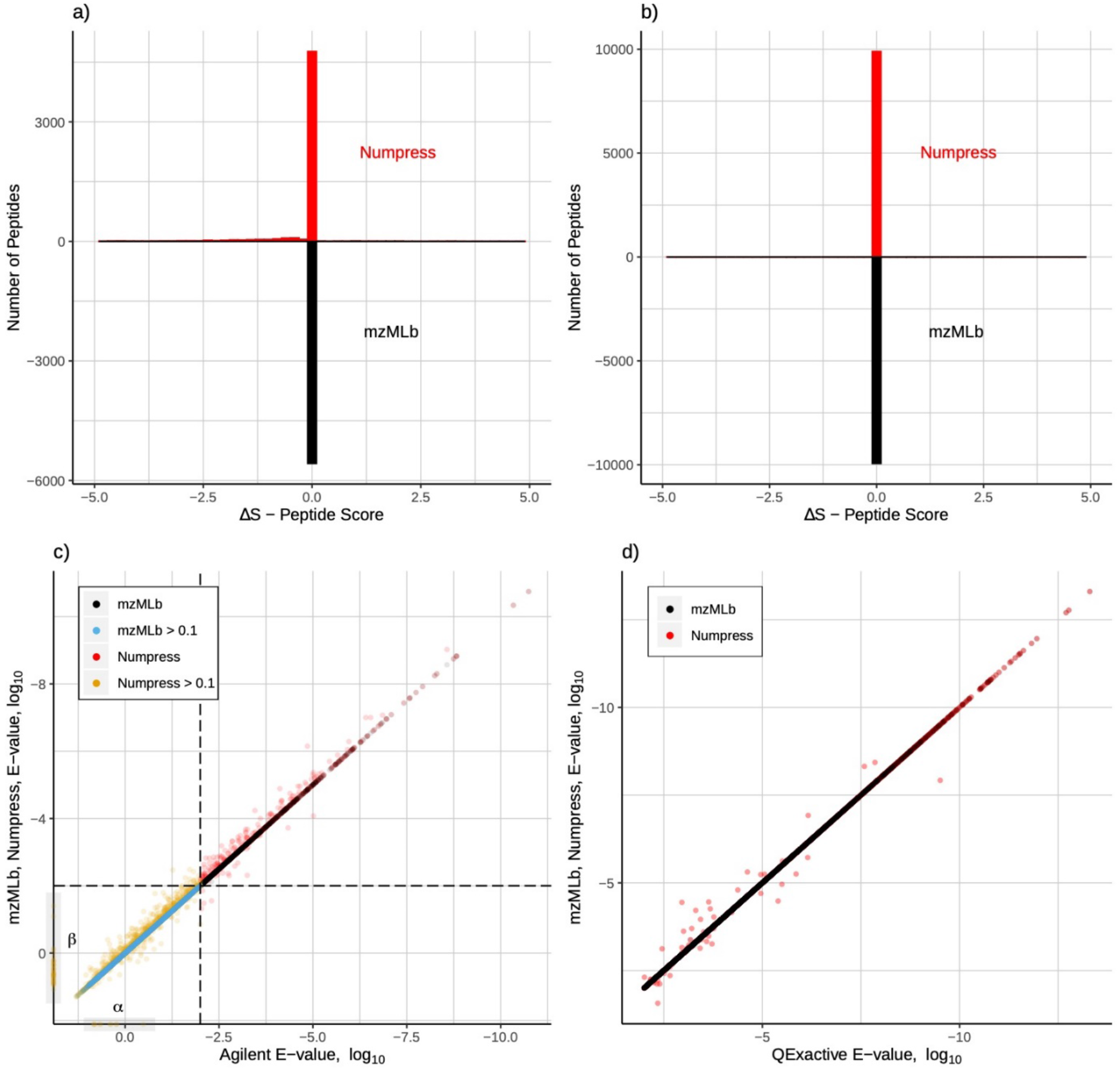
Mascot peptide PSM search results of original dataset against both Numpress and mzMLb compression for AgilentQToF and QExactive datafiles. The top two plots show the number of peptides found in Numpress and mzMLb against the original data with the x-axis representing the deviation (ΔS) of the peptide score from the original. a) for the Agilent file, here were can clearly see a number of peptide scores deviating from the original score for the Numpress case. b) the results for the QExactive file, where the number of peptides deviating in the Numpress case is much less when compared to the number of matching peptides. In both cases mzMLb outperforms Numpress and has virtually no peptides deviating from the original. The bottom two plots, show the relative E-value performance of both mzMLb and Numpress against the original dataset, with c) depicting the results for Agilent and d) for the QExactive datafile.

The HDF5 binary dataset chunk size can have a significant impact on access speed and file size. For the AgilentQToF file, Figure 6 compares mzML with zlib, mzML with Numpress + zlib, and mz5 with zlib, to mzMLb across a range of chunk sizes. Figure 6a demonstrates write performance on a Linux workstation; Dell T5810, Intel Xeon CPU E5-1650 v3, with 32GB ram and 3Tb HDD running Ubuntu Linux v18.04. In order to produce these results, we ran ProteoWizard msconvert 10 times converting the files from vendor format while recording the write duration. However, modern operating systems including the Linux kernel employ a sophisticated file and memory caching system; in order to mitigate this mechanism accelerating the multiple writes and reads of the data files being tested, we cleared the Linux memory cache after every invocation of msconvert. It can be seen for lower chunking values, mzMLb (with both mzLinear on/off) outperforms the other formats, and only starts to slow for chunk size of around 512 kb – 1024 kb. Figure 6b shows the relative compression of the files as the chunking sizes increase (again for both mzLinear on/off). It can be seen that at 1024 kb the benefit of increase chunking for compression of data quickly plateau while the writing speed deteriorates. The file sizes of mzMLb with mzLinear perform 77% better than mzML + zlib, 25% smaller than mzMLb + Numpress + zlib, and produce a 56% increase in compression when compared to mz5 + zlib.

**Figure 6.**
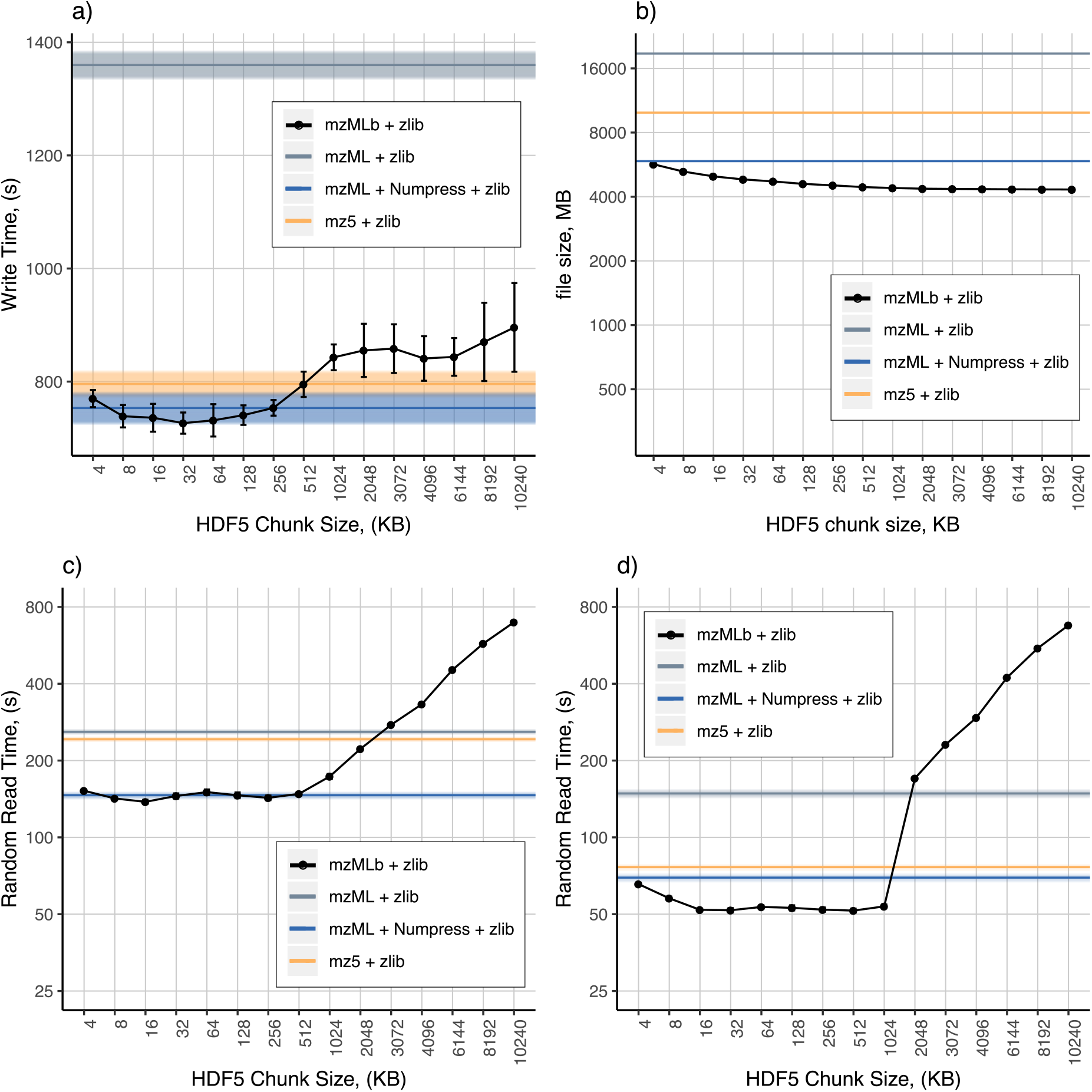
Chunk size optimization with mzLinear enabled; a) mzMLb write benchmark times with varying chunk size, b) file size with and without mzMLb enabled, c) random read benchmarks for singular spectrum access for full chunking size range, d) random read benchmarks for sequential block spectrum access for full chunking size range.

In order to evaluate the read performance of mzMLb we created a C++ program readBench, which utilizes the PreoteoWizard API and its libraries to ensure the ability to read all files formats consistently under the same software implementation. This command line tool is available from https://github.com/biospi/pwiz. Here two scenarios were considered, the first accessing a spectrum for the dataset 10,000 times at random. The second involved the random reading of 10 sequential spectra selected 1,000 times at random, thus giving 10,000 total spectrum accesses. These were also performed 10 times for each data point. The results are depicted in Figure 6c and 6d; in both cases mzMLb outperforms the other file formats while maintaining a smaller file size. Beyond 1024 kb chunk size, the random read time drastically increases.

Subsequently, we ran the random read benchmarks again but this time without zlib compression in order to evaluate use cases where fast access times are paramount and file size not important. In this test, we include mzMLb in both a lossless and lossy scenario. Here we introduced the HDF5 BLOSC^19^ plugin to the validation. The aim of BLOSC is perform modest but extremely fast decompression/compression so that the resulting read/write times are faster than using no compression at all as less data needs to be physically written to disk. It accomplishes this by: utilising a blocking technique that reduces activity on the system memory bus, transmitting data to the CPU processor cache faster than the traditional, non-compressed, direct memory fetch approach via a memcpy operating system call; leveraging SIMD instructions (SSE2 and AVX2 for modern CPUs) and multi-threading capabilities present in multi-core processors. BLOSC has a number of different optimised compression techniques including; BloscLZ, LZ4, LZ4HC, Snappy, Zlib and Zstd. Throughout these tests we used BLOSC with LZ4HC compression, as we found it to be the most effective in terms of read and write speeds when dealing with MS datasets.

In Figure 7 we depict the results of our high-throughput results designed to seek out the optimum solution for the fastest access to MS data, in two categories; lossless file formats (Figure 7a and 7b), and lossy file formats (figure 7c and 7d). In both cases we consider both random single spectra access (Figure 7a and 7c) and random block-sequential access (Figure 7b and 7d). We can see that in the case of lossless compression (Figure 7a and 7b) mzMLb performs better than both mzML and mz5 in both single and block-sequential data access.

**Figure 7.**
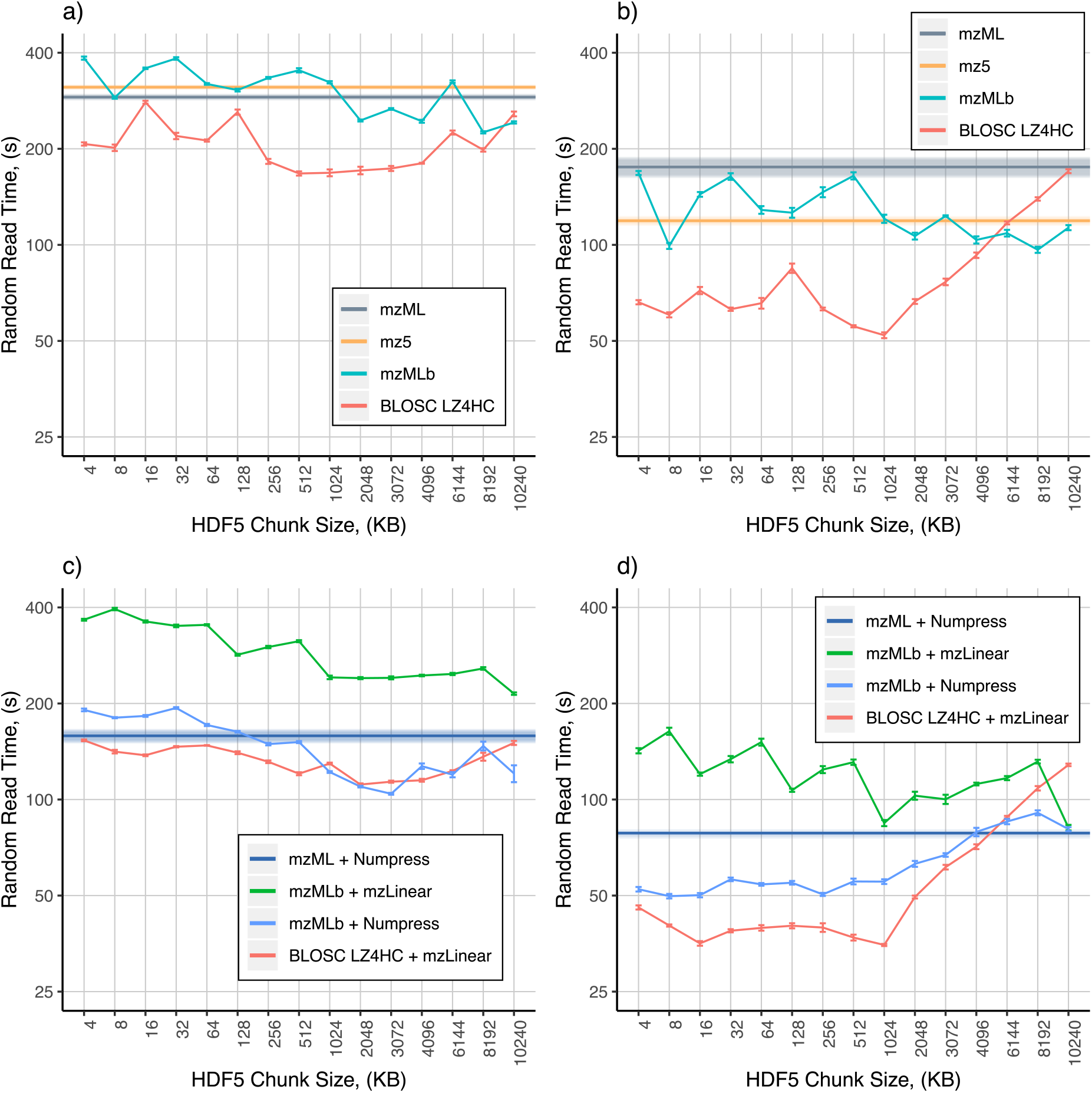
mzMLb; random read benchmarks for both; singular and block sequential, for uncompressed data with and without truncation and Numpress enabled; a) lossless single-spectrum access, b) lossless block-sequential access, c) lossy single-spectrum access, d) lossy block-sequential access.

Moreover, when we utilise mzMLb with BLOSC LZ4HC compression we can see that it significantly outperforms both mzML and mz5 at virtually all chunking sizes and particularly performs well at around 1024 kb chunks for both single and block-sequential scans. In the case of lossy datasets (figure 7c and 7d), we can see that Numpress has a significant positive impact on random read times for both single and block-sequential data access. Notably, Numpress performs better when contained within mzMLb rather than mzML. Nevertheless, when we utilize both mzMLb with mzLinear and BLOSC LZ4HC compression we observe that mzMLb is significantly faster than Numpress in block-sequential data access, and is comparable to Numpress within mzMLb for random single spectra access.

In Figure 8 and S1, we compare the file size and write performance of our new mzMLb file format against vendor raw file, mzML, mz5, and Numpress within both mzML and mzMLb. All results are the average of 10 runs. Here we also used the optimum mzMLb chunking size of 1024kb derived from both Figure 5 and Figure 6, which allows mzMLb to possess both a significant compression ratio of the file size and increased performance in both reading and writing of the mass spectrometry file. Depicted in Figure 8, the colours of the markers represent the different files formats, more specifically; the red represents mz5 files, the gray the mzML files, the blue mzMlb files and finally the orange the mzMLb files with BLOSC. The shape of the markers represents the different filters applied during the conversion process, e.g. a solid triangle represents datasets without compression, a solid diamond datasets with zlib applied, and a yellow asterisk datasets with mzLinear, truncation and zlib applied, etc. From these results it can be seen that in all cases the resulting mzMLb files were significantly smaller than mzML and a similar size to the vendor raw file. Moreover, from figure 8 and S1, mzMLb can easily be tailored to different use cases (e.g. maximum compression for archiving; lower compression but faster access times for processing, both the yellow asterisk and the solid circle markers (figure 8), representing mzLinear+trunc+zlib and Numpress+zlib respectively) in order to maximise the desired performance metric.

**Figure 8.**
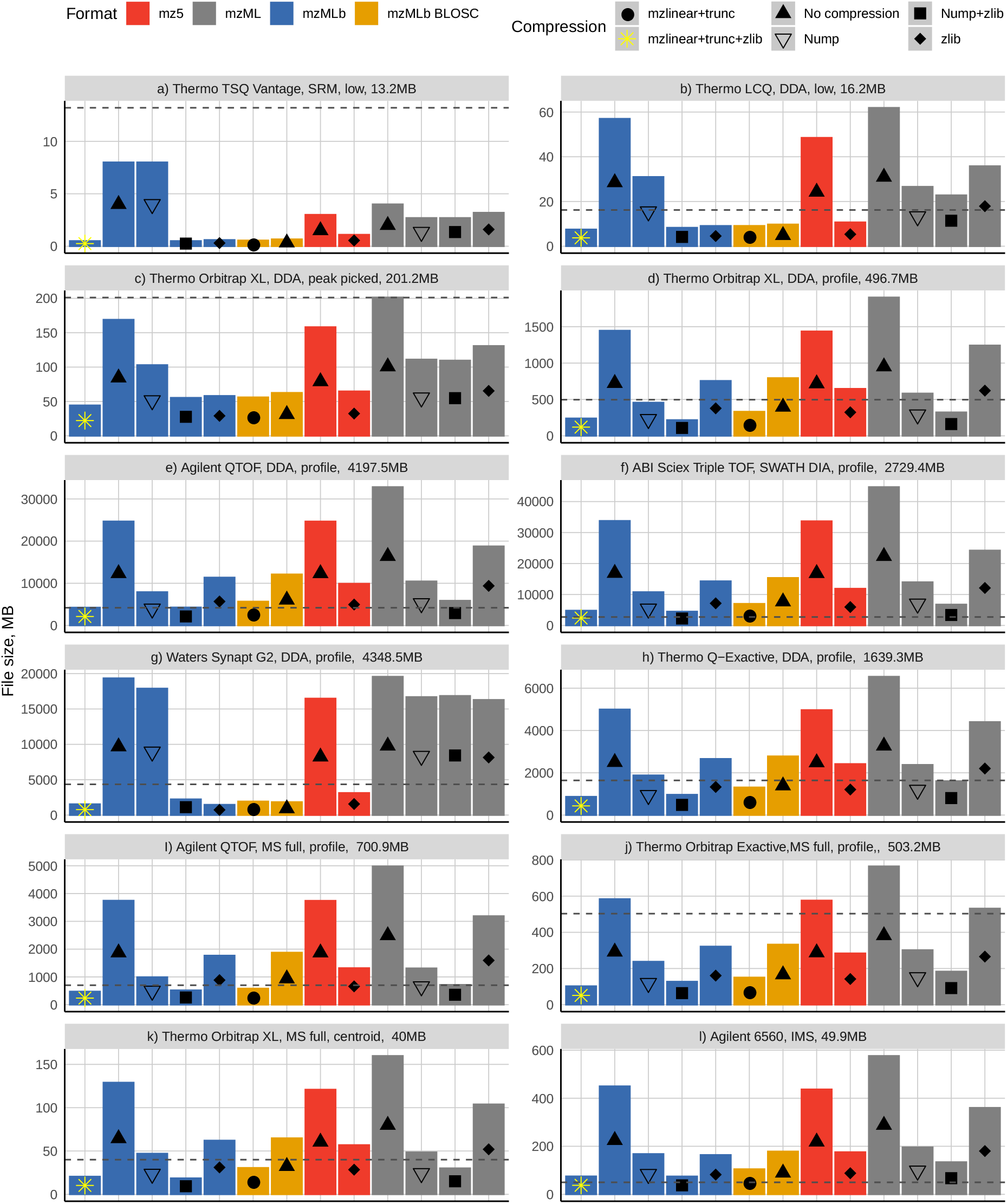
Summary data showing file sizes for all datasets using the 3 formats; mzML, mz5 and mzMLb with 6 different compression combinations spanning both lossless and lossy configurations. Uncompressed data files are also depicted here along with mzMLb BLOSC, demonstrating that fast access read times can be achieved without sacrificing file size. The original vendor file sizes are represented by the vertical dashed line.

## Conclusions

We demonstrate that by using a hybrid file format based on storing XML metadata together with native binary data within a HDF5 file it is possible to improve data reading/writing speed of raw MS data as well as preserve all related metadata in PSI-compliant mzML in an implicitly future proof way. The mzMLb file format can be tailored for different applications by changing the chunk size parameter, i.e. it is possible to adjust the format for fast access where file size does not matter, e.g. visualisation and processing, or a smaller compressed file size with slower reading/writing times for data archival. As a chunk can contain more than one spectrum of data, compression can occur across spectra, which is not possible in mzML. Our results illustrate that a chunk size of 1024kb is a good compromise for most applications

As mzMLb utilizes HDF5 we are able to leverage transparent mechanisms for random data access, caching, partial reading or writing, and error checksums, and is easily extendable through plugins to support additional filters and compression algorithms. HDF5 also allows the user to add extra information to the data file while still maintaining PSI compatibility, simply by adding extra HDF5 groups and datasets. This allows the user to store other data within the file side-by-side with the mzMLb data, for example, a version of the data optimised for fast visualisation^20^.

As we use standard features of HDF5, mzMLb is also bit-for-bit compatible with NetCDF4 (which has native Java libraries). This enables it to be easily implemented by third party processing software, as both HDF5 and NetCDF4 is widely supported across common programming languages including Java. As of v4.5.0, NetCDF also has support to allow mzMLb files to be randomly accessed remotely over the internet (the HDF5 Group have also recently delivered their own implementation of this functionality too), opening up the potential for public repositories to provide new tools for users to efficiently query and visualise their raw data archives.

## Acknowledgements

This work was supported by BBSRC grants BB/M024954 & BB/R021430, MRC grant MR/N028457 to Andrew W. Dowsey and Andrew R. Jones and BBSRC grants BB/K01997X/1 & BB/R02216X/1 to Andrew R. Jones. Eric W. Deutsch also acknowledges support from National Institutes of Health grants R01GM087221, R24GM127667, U19AG023122, and from National Science Foundation grants DBI-1933311 and IOS-1922871. We thank the Proteomics Standards Initiative community for their comments and suggestions.

## Supplementary material

**Table S1.** The raw MS data files used in the validation, together with MS instrument and method information.

| MS Data File Name | Resolution | Type | File Size (MB) | Vendor | Peak Picked | Instrument |
| --- | --- | --- | --- | --- | --- | --- |
| 101125_JT_pl3_05 | low | SRM | 13.2 | Thermo | no | TSQ Vantage |
| 110620_fract_scxB05 | low | DDA | 16.2 | Thermo | no | LCQ |
| peakPicked_121213_PhosphoMRM_TiO2_discovery | high | DDA | 201.2 | Thermo | yes | Orbitrap XL |
| 121213_PhosphoMRM_TiO2_discovery | high | DDA | 496.7 | Thermo | no | Orbitrap XL |
| AgilentQToF | high | DDA | 4197.51 | Agilent | no | QToF |
| ABSciexTripleToF | high | SWATH DIA | 2729.429 | ABI Sciex | no | Triple ToF |
| SynaptG2 | high | DDA | 4348.531 | Waters | no | Synapt G2 |
| QExactive | high | DDA | 1639.345 | Thermo | no | Q-Exactive |
| 16_PBQC-10_522-16470 | high | MS full | 700.9 | Agilent | no | QTOF |
| Orbitrap_Exacte_MS1 | high | MS full | 503.2 | Thermo | no | Orbitrap Exacte |
| Orbitrap_LTQ_MS1 | high | MS full | 40 | Thermo | no | Orbitrap XL |
| 20150813_11_pos_MM | high | Ion Mobility | 49.9 | Agilent | no | Agilent 6560 |

**Table S2.** mzMLb optimized values of the mantissa for both m/z and intensities for the data files listed in Table S1. The associated errors and files sizes for mzML with Numpress and mzMLb are also shown. Both formats used zlib compression with a compression strength of 4. For all results we show the maximum error across the whole dataset.

| MS Data File | Error |  |  |  | Truncation mzMLb |  | File Size (MB) |  |
| --- | --- | --- | --- | --- | --- | --- | --- | --- |
|  | Numpress mz | mzMLb - mz | Numpress - Intensity | mzMLb - Intensity | mz | Intensity | Numpress | mzMLb |
| 110620_fract_scxB05 | 5.49991269999423E-10 | 4.65658345231104E-10 | 0.000199901743184 | 0.000121534190427 | 31 | 10 | 22.8 | 7.5 |
| peakPicked_121213_PhosphoMRM_TiO2_discovery | 7.56648171041308E-10 | 0 | 0.000138687425465 | 0.000121936232952 | 29 | 10 | 109.6 | 44.4 |
| 121213_PhosphoMRM_TiO2_discovery | 7.56648171041308E-10 | 4.65645650512267E-10 | 0.000200016681221 | 0.000121936232952 | 31 | 10 | 323.5 | 239.9 |
| AgilentQToF | 2.63499903199379E-10 | 2.32823792996109E-10 | 0.000199380233405 | 0.000118126288434 | 32 | 10 | 5874.6 | 4248.3 |
| ABSciexTripleToF | 1.76755158654755E-09 | 9.31307141007042E-10 | 0.000149589221896 | 0.000119826764734 | 30 | 10 | 6766.2 | 4796.2 |
| SynaptG2 | 1.99996902065474E-09 | 0 | 0.000146825102953 | 0 | 29 | 10 | 16855.2 | 1545 |
| QExactive | 1.99962500899505E-09 | 1.86259363937745E-09 | 0.000200007520574 | 0.000121936232952 | 29 | 10 | 1608.4 | 868.8 |
| 16_PBQC-10_522-16470 | 2.32833492130037E-10 | 2.32811218592951E-10 | 0.000199543675631 | 0.000121042874911 | 32 | 10 | 715.5 | 475.2 |
| Orbitrap_Exacte_MS1 | 1.99502158233064E-09 | 1.86229298145969E-09 | 0.00019993678766 | 0.000121936232952 | 29 | 10 | 183.5 | 102.4 |
| Orbitrap_LTQ_MS1 | 5.77059529637374E-10 | 4.6534869138726E-10 | 0.000194229158125 | 0.000121936232952 | 31 | 10 | 30.1 | 20.5 |
| 20150813_11_pos_MM | 1.99008930234944E-09 | 1.85722806985345E-09 | 0.000163805627741 | 9.11549330011243E-05 | 29 | 10 | 134.2 | 74.9 |

**Figure S1.**
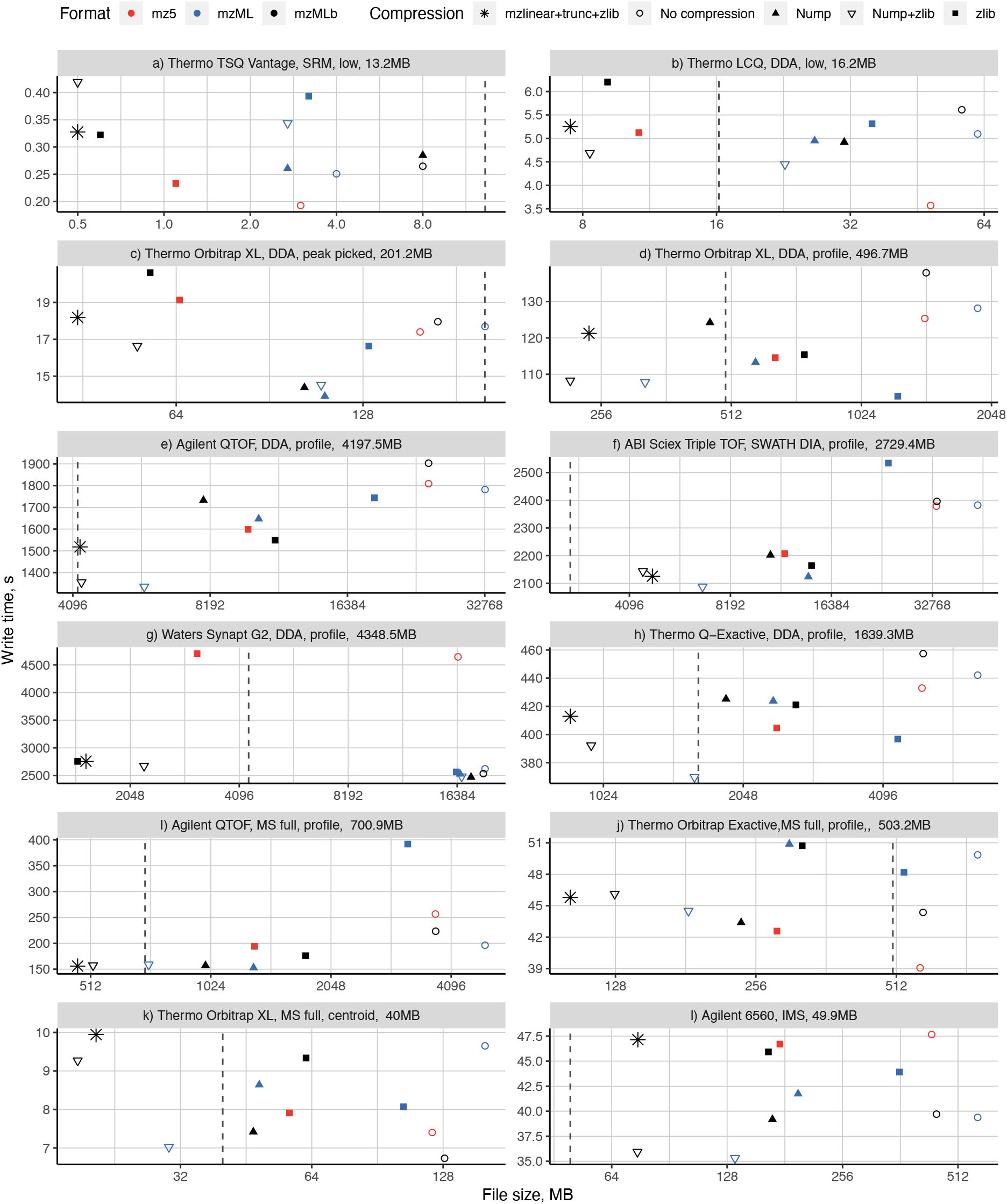
Summary data showing write times and file sizes for all datasets using the 3 formats; mzML, mz5 and mzMLb with 5 different compression combinations spanning both lossless and lossy configurations. The original vendor file sizes are represented by the vertical dashed line.

### The mzMLb format

An mzMLb dataset is a HDF5 file which must include in its root a HDF5 dataset ***mzML*** with fixed length string attribute ***version***. The currently supported version string is: “mzMLb 1.0”. The mzML XML document is stored in the ***mzML*** dataset, which is a 1D character array, with two modifications:

1. HDF5 binary indexes replace the <*indexedmzML*> wrapper schema. Here:

- HDF5 datasets ***mzML_spectrumIndex*** and ***mzML_chromatogramIndex*** replace the respective <*indexedmzML*> <*index*> blocks. Each is a 1D array of 64bit integers replicating the set of *<offset>* file pointer offsets - except note that there is an extra offset at the end of each array representing one past the end position of the last spectrum/chromatogram.
- HDF5 datasets ***mzML_spectrumIndex_idRef*** and ***mzML_chromatogramIndex_idRef***, 1D character arrays, then replicate the idRef attributes of *<offset>* as null-terminated strings concatenated together.
- Similarly and optionally, *spotID* attributes can be stored in HDF5 1D character array datasets ***mzML_spectrumIndex_spotID*** and ***mzML_chromatogramIndex_spotID***, while *scanTime* attributes can be stored in HDF5 1D floating point array dataset ***mzML_spectrumIndex_scanTime***.
2. The mzML base64 encoded binary data is removed from the <mzML> and moved into one or more native binary HDF5 datasets. Floating point binary data (*i.e.* all non-Numpress compressed *<BinaryDataArray>*) is stored as one or more HDF5 1D floating point arrays, while Numpress data can be stored as a non-base64 encoded bytestream with HDF5 data type ***OPAQUE***.

As in imzML, the mzML is modified slightly to specify this linkage to external data; the resulting XML is still valid mzML. Here, any <*binary*> blocks within the <*binaryDataArray*> blocks are removed, and the *encodedLength* attribute is set to “0”. To link to the native HDF5 binary data, within the <*binaryDataArray*> three <*cvParam*> tags need to be given, specifying the external dataset name, offset to the start of the relevant data within the dataset, and the length of the relevant data. These three tags enable flexibility over the nature and number of HDF5 datasets used to store the binary data (*e.g.* separate datasets can be used to store different datatypes; multiple spectra can be stored in the same dataset for improved chunking and compression).

### ProteoWizard mzMLb msconvert arguments

In order to convert input data into the mzMLb format using **msconvert**; the following new arguments have been introduced that allow you to alter the default parameters of converting files to mzMLb when using “**--mzMLb**” switch.

### --mzTruncation=[0-] --intenTruncation=[0-]

Perform lossy compression by removing the last n bits of mantissa from floating point data before storage. The default is 0 (no removal). Set to -1 to truncate to integers.

### --mzDelta --intenDelta --mzLinear --intenLinear

Store mz/rt or intensity values after delta or linear prediction. Predictive encoding of mz/rt values may lead to moderate improvements in gzip compression, or further improvements after floating point precision loss.

### --mzMLbChunkSize=[4096-]

Defines the chunk size to use for the mzML and all binary HDF5 datasets, in bytes. A smaller amount improves random access speed at the detriment of compression efficiency.

### --mzMLbCompressionLevel=[0-9]

Define to use either no compression (0) or GZIP compression strength 1 to 9. Compression is applied to the mzML and all binary HDF5 datasets. Specifying **--zlib** or **-z** instead will use the default compression strength of 4. If no compression is specified, the default chunk size is 1024 KB. If compression is specified, the defaults are chunk size 1024 KB, mzLinear on, mzTruncation 19 and intenTruncation 7 (as described in the main manuscript).

